# BISON: Brain tISue segmentatiON pipeline using T1-weighted magnetic resonance images and a random forests classifier

**DOI:** 10.1101/747998

**Authors:** Mahsa Dadar, D. Louis Collins

## Abstract

**Introduction:** Accurate differentiation of brain tissue types from T1-weighted magnetic resonance images (MRIs) is a critical requirement in many neuroscience and clinical applications. Accurate automated tissue segmentation is challenging due to the variabilities in the tissue intensity profiles caused by differences in scanner models and acquisition protocols, in addition to the varying age of the subjects and potential presence of pathology. In this paper, we present BISON (Brain tISue segmentatiON), a new pipeline for tissue segmentation.

**Methods:** BISON performs tissue segmentation using a random forests classifier and a set of intensity and location priors obtained based on T1-weighted images. The proposed method has been developed and cross-validated based on multi-center and multi-scanner manual labels of 72 subjects aging from 5-96 years old, ensuring the generalizability of the results to new data from various age ranges. In addition, we assessed the test-retest reliability of BISON on 2 datasets; a. using 20 subjects that had scan/re-scan MRIs and manual segmentations available, and b. using a human phantom dataset including 90 scans from a single individual acquired across 10 years.

**Results:** The results of the proposed method were compared against Atropos, a commonly used tissue classification method from ANTs. The proposed method yielded cross-validation Dice Kappa values of κ_GM_ = 0.88 ± 0.03, κ_WM_ = 0.85 ± 0.03, κ_CSF_ = 0.77 ± 0.11, outperforming ANTs Atropos (κ_GM_ = 0.79 ± 0.05, κ_WM_ = 0.84 ± 0.05, κ_CSF_ = 0.64 ± 0.22) as well as test-retest Dice Kappa values of κ_GM_ = 0.94 ± 0.006, κ_WM_ = 0.92 ± 0.006, κ_CSF_ = 0.77 ± 0.11 outperforming both manual (κ_GM_ = 0.92 ± 0.01, κ_WM_ = 0.91 ± 0.01, κ_CSF_ = 0.74 ± 0.03) and ANTs Atropos (κ_GM_ = 0.87 ± 0.001, κ_WM_ = 0.92 ± 0.001, κ_CSF_ = 0.79 ± 0.05). The human phantom dataset validations showed high generalizability for both Atropos (κ_GM_ = 0.97 ± 0.01, κ_WM_ = 0.96 ± 0.01, κ_CSF_ = 0.93 ± 0.02) and BISON (κ_GM_ = 0.95 ± 0.01, κ_WM_ = 0.94 ± 0.01, κ_CSF_ = 0.85 ± 0.03), while Atropos tended to consistently under-segment the cortical CSF. Finally, our assessment of BISON, Atropos, FAST from FSL, and SPM12 segmentations in presence of white matter hyperintensities (WMHs) showed that BISON outperforms the other three methods, correctly detecting WMHs as WM.

**Conclusion:** Our results show that BISON can provide accurate and robust segmentations in data from different age ranges and various scanner models, making it ideal for performing tissue classification in large multi-center and multi-scanner databases.

## 1. Introduction

Accurate voxel-wise segmentation of different tissue types (i.e. gray matter (GM), white matter (WM), and cerebrospinal fluid (CSF)) in magnetic resonance images (MRIs) is important in many neuroimaging applications (González-Villà et al., 2016). Regional and overall volumetric differences and changes in different tissues can be used to assess disease severity and progression. Tissue classification is also a necessary step in many image processing pipelines and applications (Ad-Dab’bagh et al., 2006; De Boer et al., 2009; Mateos-Pérez et al., 2018; Sajja et al., 2006; Schmidt et al., 2012; Simões et al., 2013; Steenwijk et al., 2013), as well as for functional activation localization in functional MRI (fMRI) processing algorithms (Jo et al., 2010). Manual tissue classification is subject to rater variability and time consuming, and therefore impractical, especially in large datasets.

Tissue segmentation is generally performed based on T1-weighted MR images, which provide a high inter-tissue contrast. However, accurate tissue segmentation can be challenging due to the presence of imaging artefacts such as noise, partial volume effects, and magnetic field non-uniformities (Cabezas et al., 2011), differences in scanner models and image acquisition protocols, as well as intensity profiles and anatomical variabilities across subjects from different ages and with different pathologies. Many researchers have attempted to address this problem, using either atlas-based techniques (Aljabar et al., 2009; Cabezas et al., 2011; Collins and Evans, 1997; Collins and Pruessner, 2010; Klein and Tourville, 2012; Lötjönen et al., 2010; Shen and Davatzikos, 2001; Wu et al., 2007) where one (or multiple) atlas with tissue (or structure) labels is nonlinearly registered to the subject volume. Using the obtained transformation, the labels of the atlas are then transformed to the subject volume. Such methods tend to provide better segmentations for the subcortical regions, but are generally not very accurate in the cortex, where there is more subject variability. Another commonly used approach is intensity-based classification, where a machine learning technique is trained on voxel-wise intensity and location features (Cocosco et al., 2003; Dadar et al., 2017a; Duda et al., 2001; Fischl et al., 2002; González-Villà et al., 2016; Makropoulos et al., 2014; Scherrer et al., 2009; Van Leemput et al., 1999). While these methods generally perform well on data with similar intensity profiles as their training libraries, they are susceptible to inaccuracies in cases where the input images have different intensity distributions (due to scanner or imaging protocol differences, age differences, or in cases of disease) from the images they have been trained on.

One of the major challenges when performing tissue classification on aging subjects or individuals with neurodegenerative diseases is the presence of white matter hyperintensities (WMHs). WMHs are areas of increased signal on T2-weighed and FLAIR MRI sequences, which appear as hypointense on T1w images (Dadar et al., 2018b). These hypointense WMH regions can present with a similar intensity profile to the gray matter in T1w images and can be segmented as GM by tissue classification methods that are solely or highly dependent on image intensities. This is particularly important in neurodegenerative disease populations (e.g. Alzheimer’s disease and Parkinson’s disease), where WMHs are very common findings (van der Flier et al., 2018) and have been shown to interact with neurodegeneration and contribute to cognitive deficits (Dadar et al., 2018c, 2019) and therefore their misclassification as GM may systematically bias the findings of such studies.

In this paper, we present BISON (Brain tISsue segmentatiON) pipeline, an intensity and location-based tissue segmentation method that has been trained and validated on Neuromorphometrics dataset, a manually segmented library of 92 subjects, aged from 5 to 96 years old, from 4 different datasets. We further validated the performance of BISON on a subset of the Neuromorphometrics dataset that had test-retest scans available and a human phantom dataset, containing 90 scans acquired across 11 different sites to ensure the robustness and generalizability of the results. This multi-center and multi-scanner training and validation ensure the generalizability of the results to data from different scanners and age ranges. In addition, we make the segmentation pipeline along with the pre-trained classifier publicly available (http://nist.mni.mcgill.ca/?p=2148).

## 2. Methods

### 2.1. Data

Data used in this paper was obtained from the Neuromorphometrics database of neuroanatomically labelled MRI brain scans (http://www.neuromorphometrics.com/?page_id=23#demo) (Caviness Jr et al., 1999). The database includes 92 unique subjects from 4 different studies, namely, the Alzheimer’s Disease <u>Neuroimaging</u> Initiative (ADNI) database (adni.loni.usc.edu), the 20 Repeats dataset of 20 nondemented subjects scanned on two visits within 90 days and labelled by two raters, the Child and Adolescent NeuroDevelopment Initiative (CANDI) database (http://www.nitrc.org/projects/candi_share), and the Open Access Series of Imaging Studies (OASIS) database (http://www.oasis-brains.org/). Tale 1 shows a summary of the demographic information for each data set.

#### Human Phantom Dataset

A second dataset of 90 scans acquired from one subject (healthy male, aged 42-51 years) between years 2008 and 2017 across 11 different sites was used to further assess the generalizability of the segmentations to data from different scanners.

### 2.2. MR Images

Table 2 summarizes the acquisition parameters for each dataset included in Noromorphometrics as well as the range of the parameters in the Human Phantom dataset.

**Table 1.** Demographic information for ADNI 30, 20 Repeats, CANDI 13, and OASIS 30 datasets.

| Dataset | Min Age | Max Age | No. of Scans | No. of Subjects | Repeat Scans |
| --- | --- | --- | --- | --- | --- |
| ADNI 30 | 62.4 | 87.9 | 30 | 29 | 1 |
| 20 Repeats | 19 | 34 | 40 | 20 | 20 |
| CANDI 13 | 5 | 15 | 13 | 13 | 0 |
| OASIS 30 | 18 | 96 | 30 | 30 | 0 |

**Table 2.**
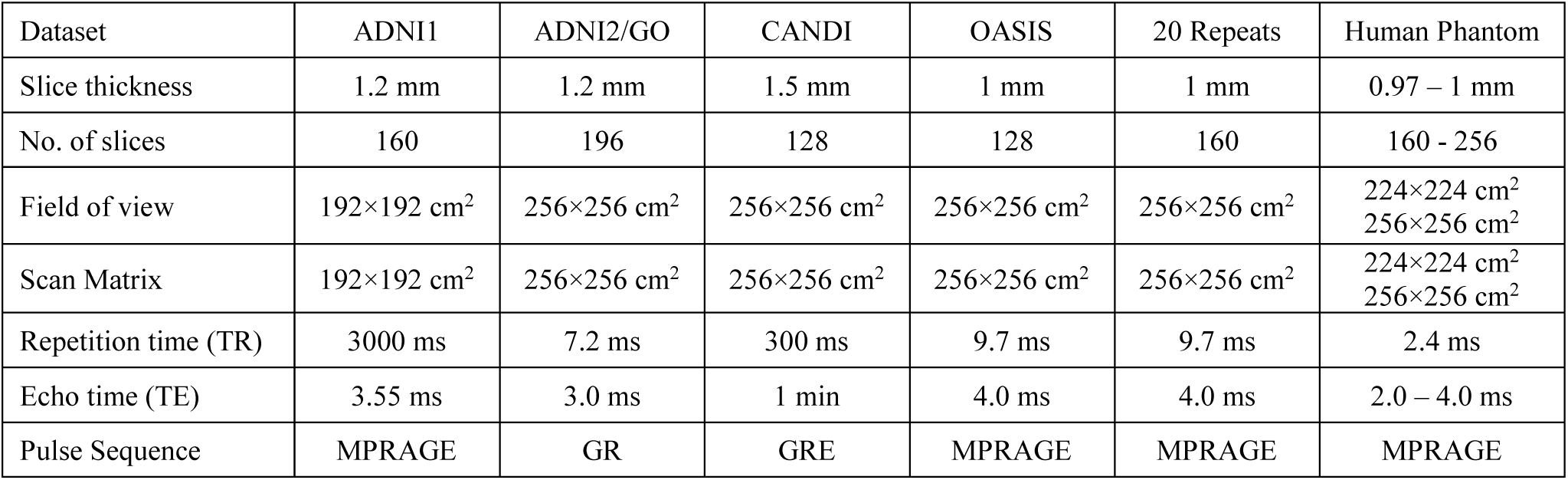
Scanner information and MRI acquisition parameters for Neuromorphometrics and Human Phantom datasets.

### 2.3. Manual Labels

The manually segmented labels were created and edited by highly trained neuroanatomical technicians using Neuromorphometrics software (Worth et al., 2001). This tool allows the use of intensity histograms, iso-intensity contours and the manual drawing and erasing of borders where necessary, allowing the user to efficiently delineate and label anatomy in 3 dimensions. The segmentation protocol specifications of Neuromorphometrics dataset can be found in http://Neuromorphometrics.com/ParcellationProtocol_2010-04-05.PDF.

### 2.4. Preprocessing

All MRI images were preprocessed using MINC toolkit, publicly available at https://github.com/BIC-MNI/minc-tools through the following steps: (I) denoising (Coupe et al., 2008), (II) intensity non-uniformity correction (Sled et al., 1998), and (III) image intensity normalization into range (0-100) using a linear intensity histogram matching algorithm. All T1-weighted images were both linearly and non-linearly registered to the MNI-ICBM152 template (Collins and Evans, 1997; Dadar et al., 2018a) to enable the use of anatomical priors in the segmentation.

To assess the test-retest reliability of the proposed method, the two T1-weighted scans for the subjects from the 20 Repeats dataset were co-registered using a 6-parameter rigid body registration. Using this transformation, the manual and automated segmentations of the second repeat were then co-registered to the first repeat for each subject.

### 2.5. Tissue Segmentation

For a given voxel with intensity value *I* from image *T* at location (*x, y, z*), the following set of intensity and location features are used to train a Random Forests classifier (Breiman, 2001) to perform tissue segmentation:

1. Voxel intensity of the preprocessed native T1-weighted image at the specific voxel location; i.e. *T*(*x, y, z*).
2. The voxel intensity of the brain template for the specific voxel location; i.e. *MNI*(*x, y, z*). To achieve this, the MNI-ICBM152 average template was nonlinearly registered to the native T1-weighted image (using the linear + nonlinear transformations).
3. Three features representing the tissue probability of the atlas for each tissue type at the specific voxel location; i.e. *P*_*GM*_(*x, y, z*), *P*_*WM*_(*x, y, z*), *P*_*CSF*_(*x, y, z*). To achieve this, the probabilistic gray matter (GM), white matter (WM), and cerebrospinal fluid (CSF) maps corresponding to the MNI-ICBM152 template were nonlinearly registered to the native T1-weighted image.
4. Three features representing the probability of the voxel belonging to each of the tissue types based on its intensity, obtained by creating probability density functions based on the intensity histograms of the tissues at all voxel locations (*PDF*_*GM*_, *PDF*_*WM*_, *PDF*_*CSF*_) from the manually labelled images in the training library; i.e. *PDF*_*GM*_(*I*), *PDF*_*CSF*_(*I*), and *PDF*_*CSF*_(*I*). *I* is the intensity value of image *T* after intensity normalization (0 < *I* < 100) at voxel location (*x, y, z*). The *PDFs* are calculated within the cross-validation loop, to avoid any possibility of *leakage (or double-dipping)* (Mateos-Pérez et al., 2018).

The Scikit-learn Python library implementation of the Random Forest classifier with 100 estimators was used (Pedregosa et al., 2011). Training and segmentations were performed in the native space of the T1-weighted images, to avoid any blurring caused by resampling of the images that might reduce the contrast at the tissue borders.

A set of 72 subjects from ADNI 30 (N=29), CANDI 13 (N=13), and OASIS 30 (N=30) were used to train the random forests classifier. Ten-fold cross validation across subjects was used to train and validate the performance of the classifier; i.e. no voxels from the subjects used for validation were used in the training stage. To avoid any overfitting caused by using multiple scans from the same subject, neither the repeat scan from the ADNI 30 subject or any of the data from 20 Repeats dataset were used in the training library. This enabled us to use the 20 Repeat dataset as a held-out validation dataset.

Dice Kappa similarity index (Dice, 1945) and volumetric correlations were used to compare the agreement between manual and automatic segmentations.

### 2.6. Comparison with ANTs Atropos

The results of the proposed method were also compared against ANTs Atropos, an ITK-based open source multi-class brain segmentation technique distributed with (ANTs http://www.picsl.upenn.edu/ANTs). Atropos performs tissue classification within a Bayesian framework by modeling the class intensities based on either parametric or non-parametric finite mixtures (Avants et al., 2011). Atropos incorporates the template-based tissue probability maps as prior information in the form of Markov Random Fields (MRF). Based on Atropos requirements, the preprocessed T1-weighted images were linearly registered to the MNI-ICBM152 template. Using the nonlinear transformations, the MNI-ICBM152 tissue probability maps were also registered to these images to be used as MRF tissue priors. Atropos was then run with default parameters to segment 3 classes on all Neuromorphometrics scans. Using the linear transformations, all the automated Atropos segmentations were then transformed back to the native space of the T1-weighted images and were compared against manual segmentations and BISON. To ensure that this resampling does not unfairly misrepresent Atropos performance, the Dice Kappas for Atropos were also calculated without this resampling step.

### 2.7. Data and Code Availability Statement

The full script for the BISON segmentation pipeline along with the random forest classifier pre-trained on the Neuromorphometrics dataset is publicly available at (http://nist.mni.mcgill.ca/?p=2148).

## 3. Results

### 3.1. Cross Validation Performance

The performance of BISON was first validated through 10-fold cross validation. All voxels within the manually segmented masks (i.e. all voxels labeled as GM, WM, or CSF by the raters) of the subjects were classified and used for validation. Figure 1 shows the boxplot of the Dice Kappa for each tissue and each dataset, for BISON and Atropos.

**Fig. 1.**
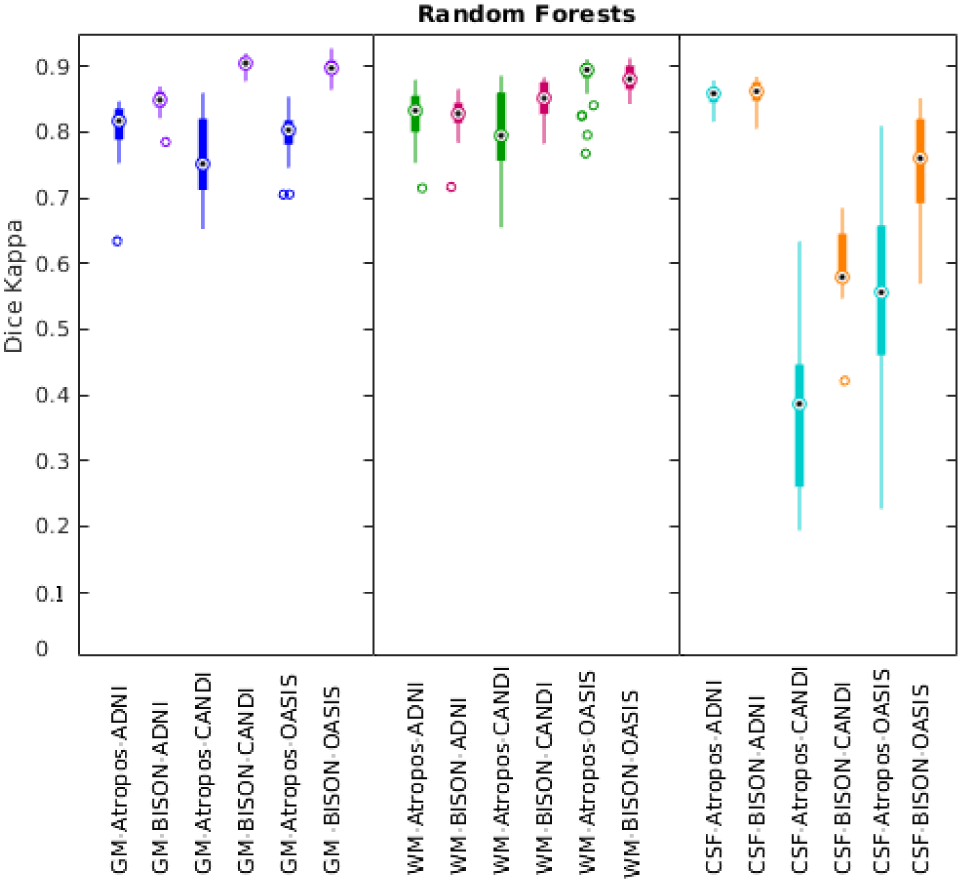
Dice Kappa plots showing the agreement between manual and automated segmentations for GM, WM, and CSF separately for ADNI, CANDI, and OASIS subjects. Circles show the median values. GM= Gray Matter. WM= White Matter. CSF= CerebroSpinal Fluid.

Overall, BISON achieved significantly higher Dice Kappa values for GM and CSF (paired t-test, p<0.0001), and marginally significant values for WM (paired t-test, p=0.05). The overall average Dice Kappa value for BISON was κ_GM_ = 0.88 ± 0.03, κ_WM_ = 0.85 ± 0.03, κ_CSF_ = 0.77 ± 0.11 and κ_GM_ = 0.85 ± 0.02, κ_WM_ = 0.83 ± 0.03, κ_CSF_ = 0.86 ± 0.02 for the ADNI subjects, κ_GM_ = 0.90 ± 0.01, κ_WM_ = 0.85 ± 0.03, κ_CSF_ = 0.59 ± 0.07 for the CANDI subjects, and κ_GM_ = 0.90 ± 0.01, κ_WM_ = 0.88 ± 0.02, κ_CSF_ = 0.74 ± 0.08 for the OASIS subjects. In comparison, ANTs Atropos obtained overall mean Dice Kappas of κ_GM_ = 0.79 ± 0.05, κ_WM_ = 0.84 ± 0.05, κ_CSF_ = 0.64 ± 0.22 and κ_GM_ = 0.81 ± 0.04, κ_WM_ = 0.82 ± 0.04, κ_CSF_ = 0.85 ± 0.02 for the ADNI subjects, κ_GM_ = 0.76 ± 0.07, κ_WM_ = 0.80 ± 0.07, κ_CSF_ = 0.38 ± 0.14 for the CANDI subjects, and κ_GM_ = 0.80 ± 0.04, κ_WM_ = 0.88 ± 0.04, κ_CSF_ = 0.55± 0.17 for the OASIS subjects. Recalculating the Dice Kappas in the MNI-ICBM152 space (without resampling back to the native T1w space) did not change Atropos results (overall mean Dice Kappa: κ_GM_ = 0.79 ± 0.05, κ_WM_ = 0.84 ± 0.05, κ_CSF_ = 0.64 ± 0.22).

The overall correlation between manual and automatic volumes was r_GM_ = 0.95, p_GM_ <0.0001, r_WM_ = 0.89, p_WM_ <0.0001, and r_CSF_ = 0.97, p_CSF_ <0.0001 for the proposed method, and r_GM_ = 0.87, p_GM_ <0.0001, r_WM_ = 0.89, p_WM_ <0.0001, and r_CSF_ = 0.70, p_CSF_ <0.0001 for ANTs Atropos. Figure 2 shows the volumetric correspondence between the manual and automatic volumes.

**Fig 2.**
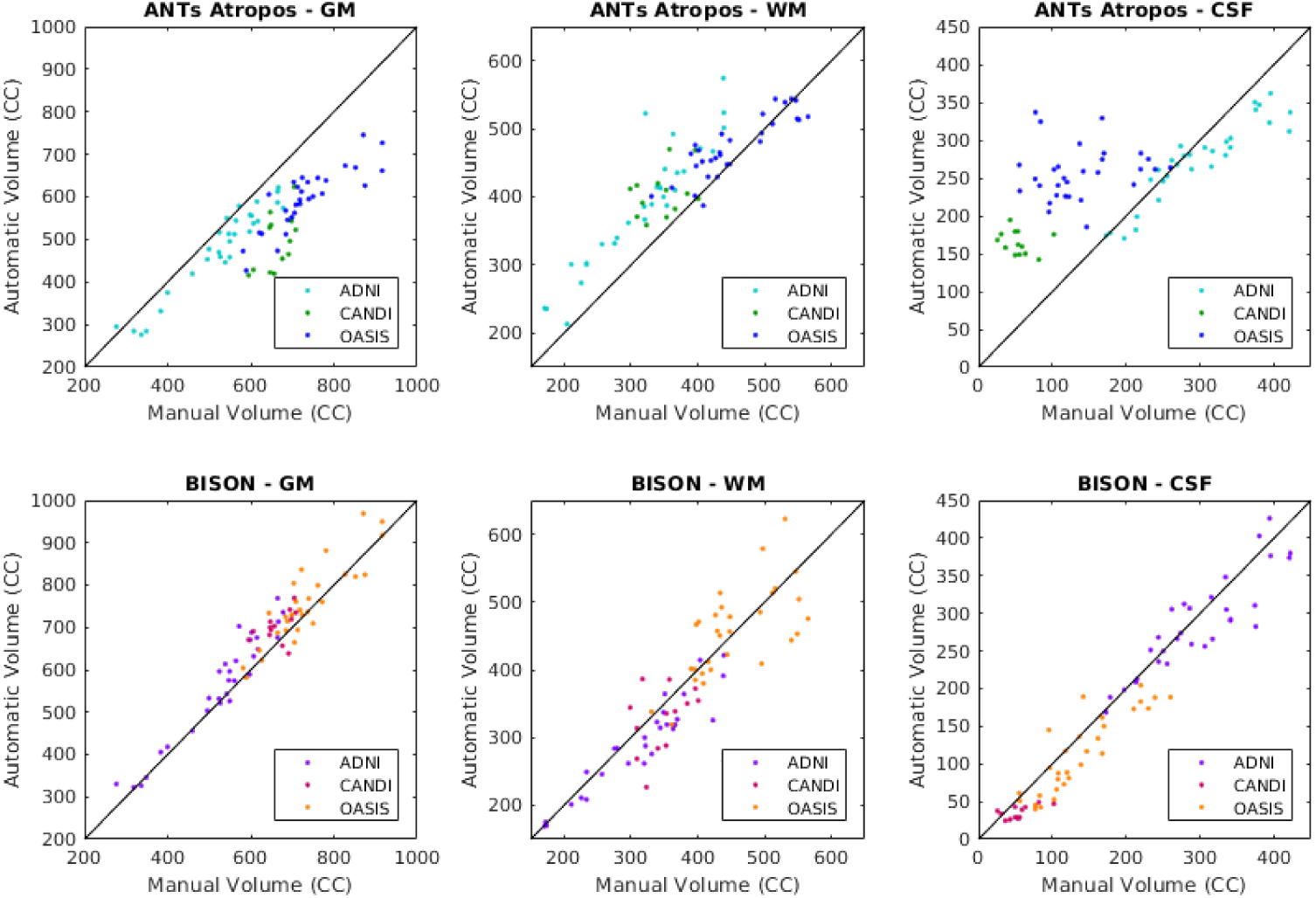
Automatic versus manual volumes for GM, WM, and CSF in ADNI, CANDI, and OASIS for the RF classifier and ANTs Atropos. GM= Gray Matter. WM= White Matter. CSF= CerebroSpinal Fluid.

Figure 3 compares the manual and automated segmentation results for one subject from each dataset.

**Fig. 3.**
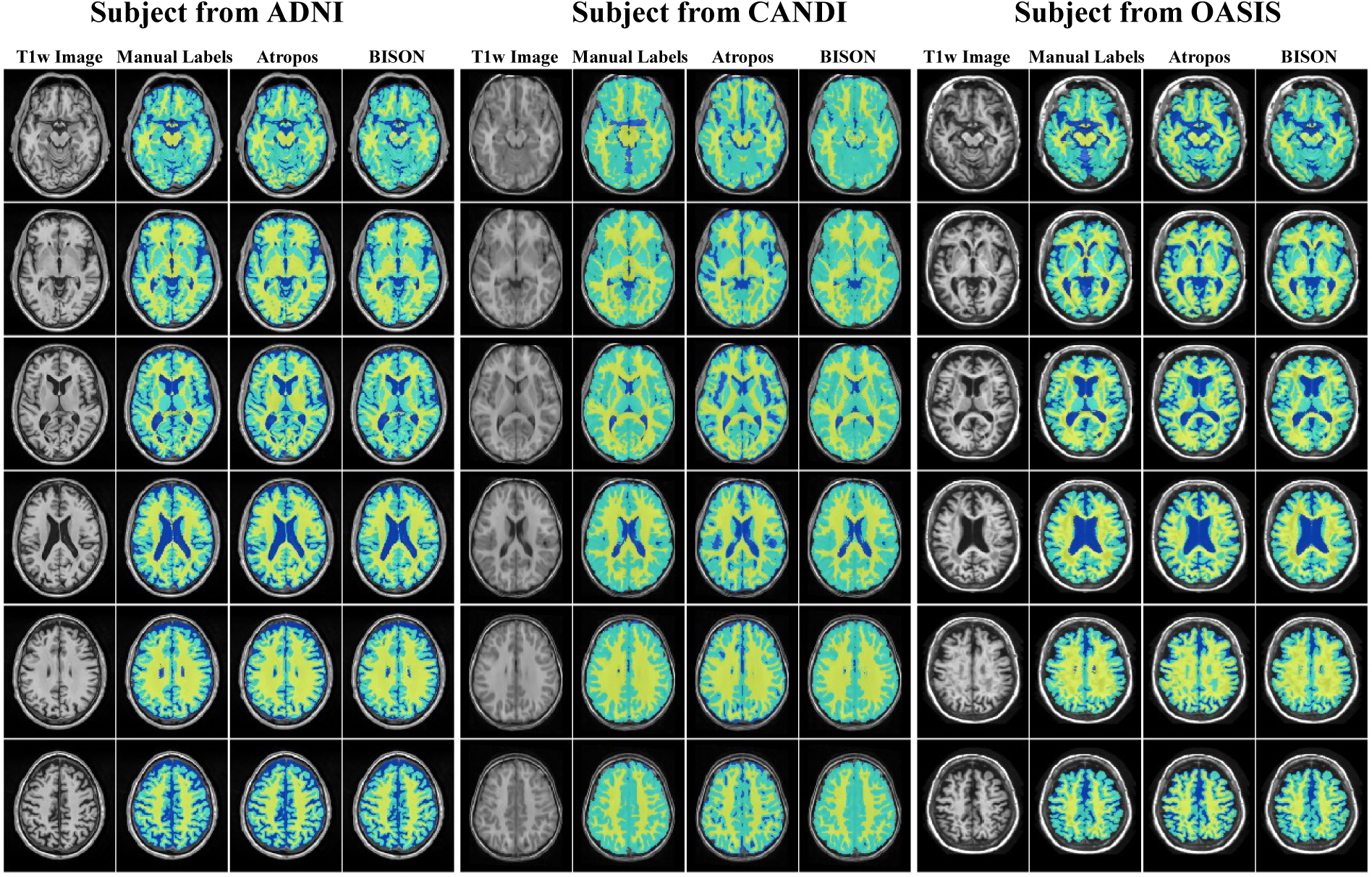
Axial slices comparing manual segmentations and automatic segmentations from ADNI, CANDI, and OASIS datasets.

### 3.2. Test-Retest Performance

Figure 4 shows the test-retest boxplots of the Dice kappa values for each tissue for both for the manually segmented labels and the automatically segmented results.

**Fig. 4.**
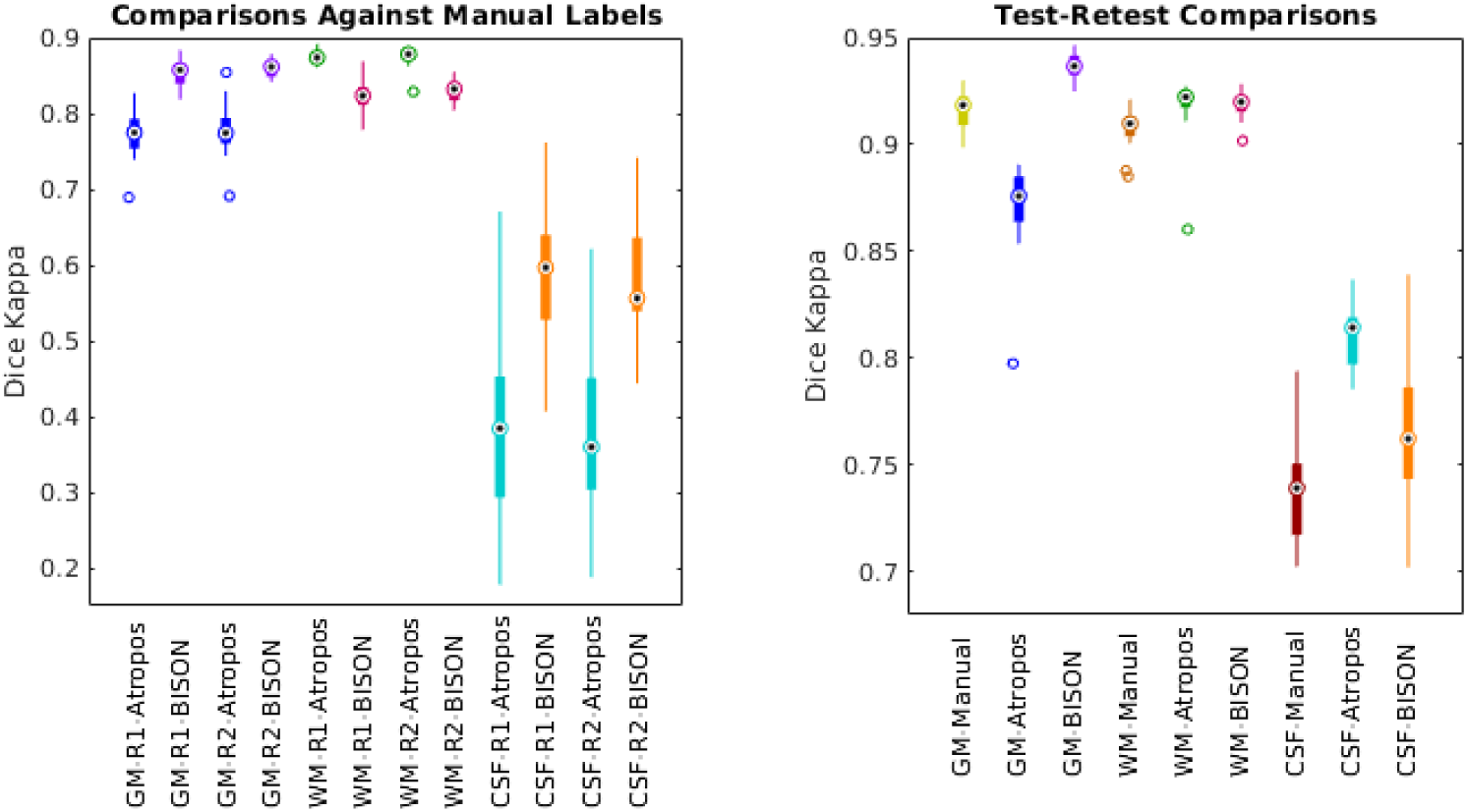
Dice Kappa plots showing the comparisons with the manual segmentations as well as the test-retest agreement for manual and automated segmentations for GM, WM, and CSF separately for 20 Repeat subjects. GM= Gray Matter. WM= White Matter. CSF= CerebroSpinal Fluid.

When comparing to manual labels of the test-retest (R1 and R2) data, the average Dice Kappa value for BISON was κ_GM_ = 0.86 ± 0.02, κ_WM_ = 0.82 ± 0.02, κ_CSF_ = 0.58 ± 0.09 for the first scan (R1), and κ_GM_ = 0.86 ± 0.01, κ_WM_ = 0.83 ± 0.02, κ_CSF_ = 0.58 ± 0.08 for the second scan (R2). In comparison, Atropos obtained mean Dice Kappas of κ_GM_ = 0.77 ± 0.03, κ_WM_ = 0.87 ± 0.01, κ_CSF_ = 0.38 ± 0.12 for the first scan, and κ_GM_ = 0.78 ± 0.03, κ_WM_ = 0.87 ± 0.02, κ_CSF_ = 0.37 ± 0.12 for the second scan. All the differences are statistically significant (paired t-test, p<0.0001).

Consistency (or reproducibility) was estimated by comparing the segmentations of test data to the retest data. For the manual segmentations, this gives the inter-rater reliability. The average test-retest Dice Kappa values were κ_GM_ = 0.92 ± 0.01, κ_WM_ = 0.91 ± 0.01, κ_CSF_ = 0.74 ± 0.03 for the manual segmentations; κ_GM_ = 0.94 ± 0.006, κ_WM_ = 0.92 ± 0.006, κ_CSF_ = 0.77 ± 0.11 for the proposed method; and κ_GM_ = 0.87 ± 0.001, κ_WM_ = 0.92 ± 0.001, κ_CSF_ = 0.79 ± 0.10 for ANTs Atropos. The differences between Atropos and BISON were statistically significant for GM (paired t-test, p<0.0001), but not for WM and CSF. Figure 5 shows the manual and automated segmentation results for both repeats for one subject. Overall, Atropos tends to over-segment CSF, and has failed to detect the putamen and thalamus in some slices, where the contrast between GM and WM is relatively low (e.g. row 3).

**Fig. 5.**
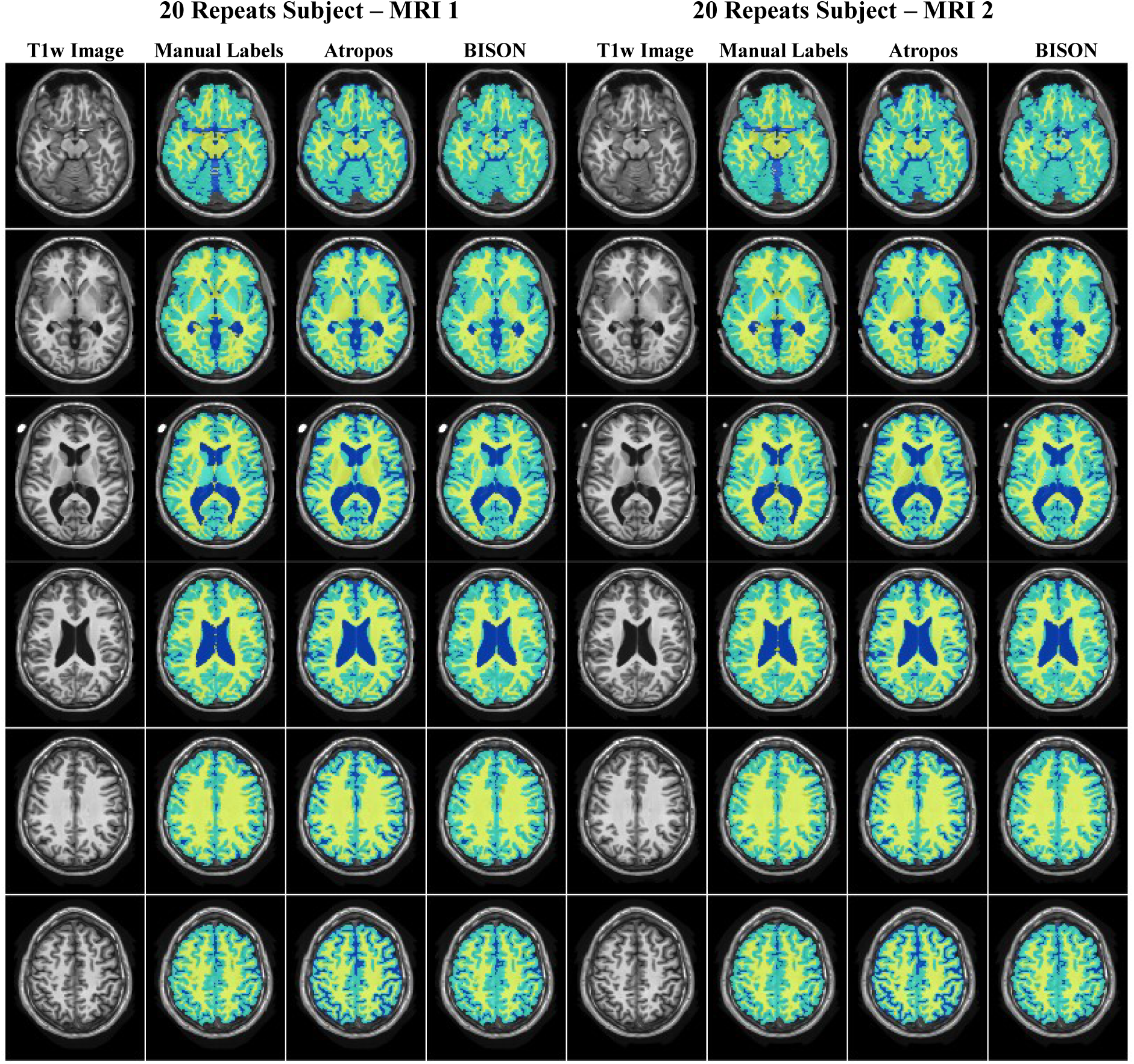
Axial slices comparing test-retest manual and automatic segmentations for a subject from 20 Repeats dataset.

### 3.3. Generalizability

In order to assess the consistency of the segmentations on data from different scanners, we segmented 90 T1w MRI scans obtained across 10 years from one subject. Figure 6 shows the volumes (normalized by the total intracranial volume) estimated based on Atropos and BISON segmentations, across 10 years. Both methods have very consistent results across different scans, with variabilities (normalized standard deviation) of 0.3%, 0.3%, 0.1% for GM, WM, and CSF for Atropos, and 0.3%, 0.2%, 0.3% for BISON, respectively. Based on Atropos results, GM volume decreases significantly (0.05% per year), while WM (0.03% per year) and CSF (0.02% per year) volumes significantly increase. BISON estimates of the tissue volumes show more modest changes in the volumes, with a slight but non-significant decrease for GM (0.01% per year) and WM (0.01% per year), and a non-significant increase in the CSF (0.02% per year) volume.

**Fig 6.**
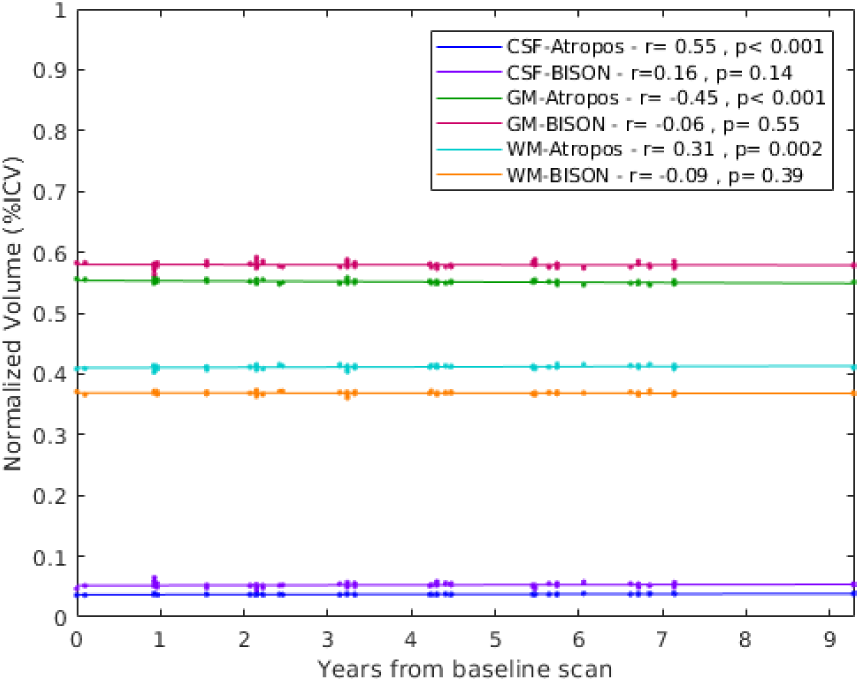
Human Phantom GM, WM, and CSF volumes obtained from 90 scans. r, p in the figure legends show the correlation between time from baseline and the volumes. GM= Gray Matter. WM= White Matter. CSF= CerebroSpinal Fluid.

Figure 7 shows the Dice Kappa values for each pair of segmentations for Atropos and BISON. The average Dice Kappa for all pairs was κ_GM_ = 0.97 ± 0.01, κ_WM_ = 0.96 ± 0.01, κ_CSF_ = 0.93 ± 0.02 for Atropos, and κ_GM_ = 0.95 ± 0.01, κ_WM_ = 0.94 ± 0.01, κ_CSF_ = 0.85 ± 0.03 for BISON, showing excellent consistency between different segmentations.

**Fig. 7.**
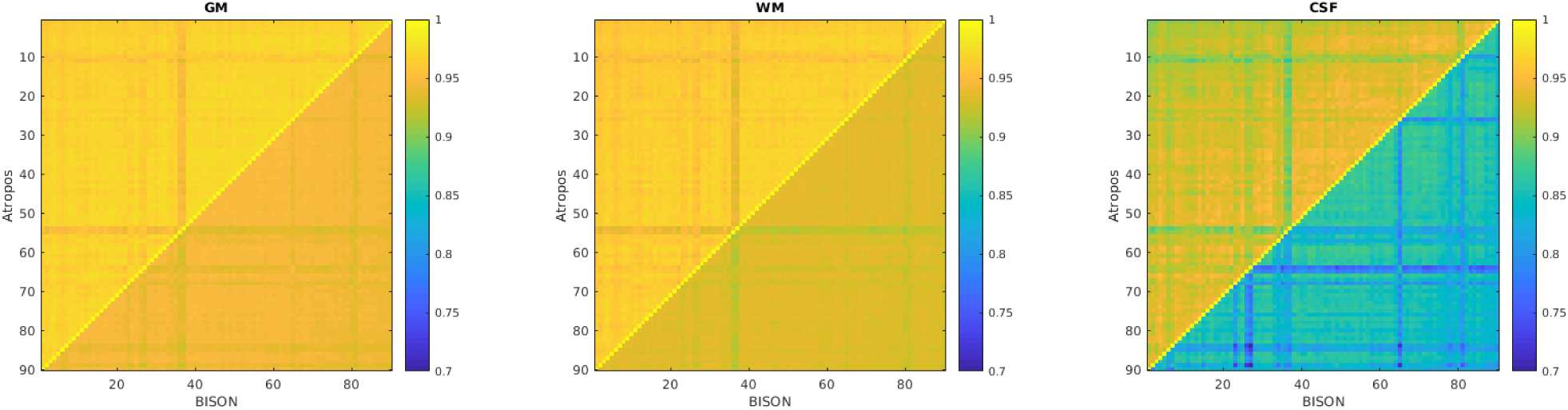
Dice Kappa values for GM, WM, and CSF, showing the agreement between each pair of scans in the Human Phantom dataset. Upper and lower triangles show Atropos and BISON results respectively. GM= Gray Matter. WM= White Matter. CSF= CerebroSpinal Fluid.

Figure 8 shows the segmentations by Atropos and BISON for one timepoint. Overall, Atropos tends to consistently under-segment the CSF in the sulci, missing a large portion of the CSF in the cortical regions (rows 5 and 6 in Figure 8, also reflected in the lower CSF volumes in Figure 6).

**Fig. 8.**
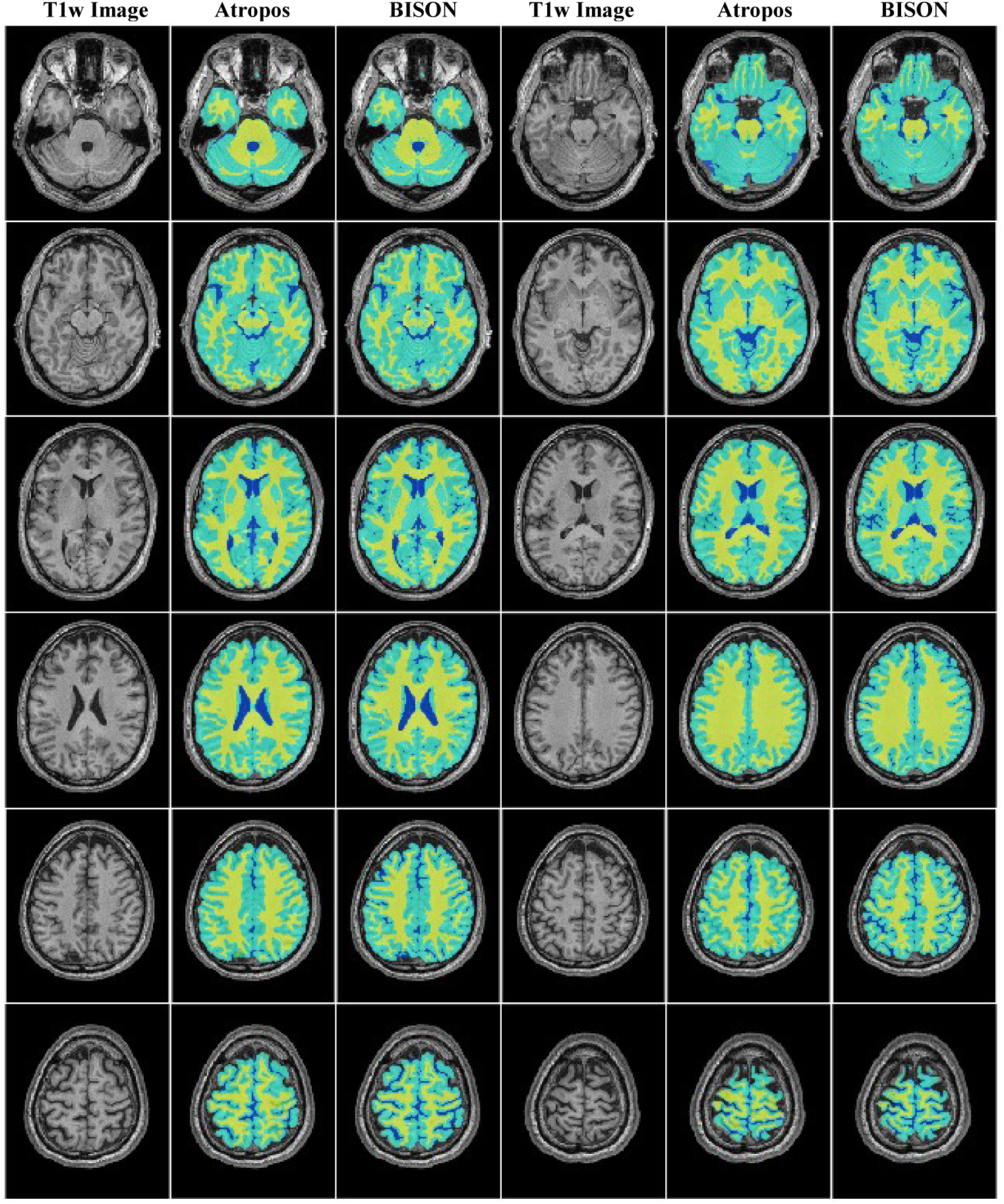
Axial slices comparing Atropos and BISON segmentations for one timepoint from the Human Phantom dataset.

### 3.4. Tissue Segmentation in Presence of Vascular Pathology

To compare the performance of BISON with other commonly used tissue classification pipelines in presence of vascular pathology, we assessed the subjects with high loads of WMHs in the Neuromorphmetrics dataset. 16 subjects with moderate to severe levels of WMHs were identified based on visual assessment. T1w images of these subjects were also segmented by SPM12 (Penny et al., 2011), and FAST from FSL (Zhang et al., 2001), in addition to Atropos and BISON. Visual qualitative assessment of the segmentation results showed that while BISON was able to successfully detect most of the WMHs as WM, Atropos and FAST segmented them as GM. SPM12 segmented most of the WMHs as WM, however, the presence of WMHs led to gross under-segmentation of the GM (and a corresponding over-segmentation of WM). Figure 9 shows axial slices covering the brain of one subject with WMHs. The WMH regions are shown with red arrows. Table 3 shows the Kappa values for GM, WM, and CSF for this subject as well as the average values for all 16 subjects for each method.

**Table 3.** Dice Kappa Values for GM, WM, and CSF for Atropos, BISON, FAST, and SPM12 for one subject with WMHs (Figure 9) as well as mean ± standard deviation of 16 subjects with WMHs. GM= Gray Matter. WM= White Matter. CSF= Cerebrospinal Fluid.

|  | Tissue Type | Atropos | BISON | FAST | SPM12 |
| --- | --- | --- | --- | --- | --- |
| Example | GM | 0.70 | 0.87 | 0.69 | 0.65 |
|  | WM | 0.76 | 0.86 | 0.81 | 0.77 |
|  | CSF | 0.55 | 0.70 | 0.47 | 0.55 |
| Average | GM | $0.79 \pm 0.04$ | $0.86 \pm 0.02$ | $0.74 \pm 0.08$ | $0.80 \pm 0.07$ |
| | WM | $0.82 \pm 0.04$ | $0.84 \pm 0.02$ | $0.82 \pm 0.08$ | $0.84 \pm 0.06$ |
| | CSF | $0.79 \pm 0.09$ | $0.84 \pm 0.04$ | $0.66 \pm 0.16$ | $0.76 \pm 0.13$ |

**Fig. 9.**
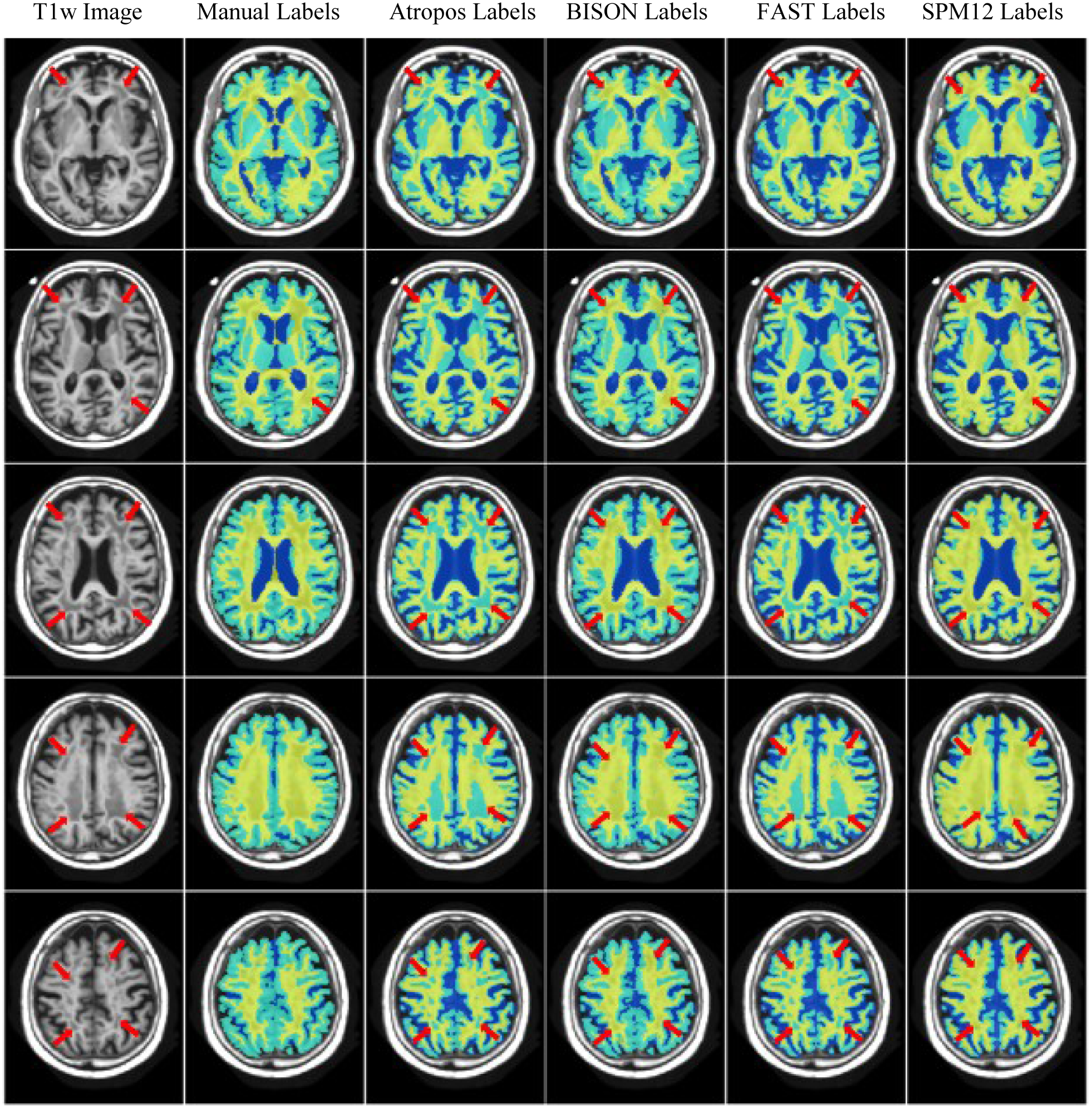
Axial slices covering the brain comparing manual segmentations and automatic segmentations from Atropos, BISON, FAST, and SPM12 for a subject with WMHs. WMH regions are indicated by red arrows. WMH= White Matter Hyperintensities.

## 4. Discussion

In this paper, we presented a robust pipeline for tissue segmentation and validated its performance in a multi-cohort and multi-scanner scanner dataset of subjects with different age ranges. The proposed method uses a set of intensity and spatial probability features as well as a Random Forest classifier to perform tissue classification. We quantitatively compared the performance of the proposed technique against ANTs Atropos in Neuromorphometrics and Human Phantom datasets, both of which include multi-center and multi-scanner images. We also compared the performance of BISON in presence of WMHs with Atropos, SPM12, and FAST from FSL; three frequently used publicly available tissue classification methods.

Training and validation of the proposed method on multi-center and multi-scanner data ensures its generalizability to new data, since automated methods that are trained based on single scanner datasets generally tend to over-specialize on the characteristics of that specific dataset (e.g. scanner model, acquisition sequence, population under study) and perform worse when applied to data from other scanners or populations (Dadar et al., 2017b). This is even more true when the automated methods are trained and validated only against synthetically generated images. Our previous experience has shown that including multi-scanner datasets in the training library can increase the generalizability of the segmentation method (Dadar et al., 2017a).

ANTs Atropos Dice Kappa values were lower than the original results reported in the paper (κ_GM_ = 0.95, κ_WM_ = 0.96, and κ_CSF_ = 0.94), however, those results were developed and validated based on synthetic data and automatically generated labels (BrainWeb 20-subject dataset: https://brainweb.bic.mni.mcgill.ca/brainweb/anatomic_normal_20.html.) of normal young brains generated by an MRI simulator (Aubert-Broche et al., 2006), which is a much less challenging problem, and also not affected by inter-rater and intra-rater variability caused by manual segmentations (Avants et al., 2011). To ensure that the lower performance was not caused by any differences in our application of Atropos, we applied Atropos to the BrainWeb dataset as well as the example provided by ANTs and were able to obtain similar results.

Atropos tended to segment CSF more generously in subjects that had lower CSF volumes (Figures 2, 3), and under-segment them in subjects with large levels of CSF. The volumetric comparisons (Figure 2) showed that Atropos generally tends to over-segment CSF and under-segment GM in CANDI (children from 5 to 15 years old with low CSF volumes) and OASIS datasets, whereas the volumes obtained by the proposed method have a linear relationship with the manual volumes.

Our experiments with the Human Phantom dataset showed excellent consistency across different scans for both Atropos and BISON, with variabilities (normalized standard deviation) of 0.3%, 0.3%, 0.1% for GM, WM, and CSF for Atropos, and 0.3%, 0.2%, 0.3% for BISON, respectively. Part of this variability comes from the fact that these scans were acquired across 10 years, while the subject aged from 42 to 51 years old, and therefore may reflect actual change in tissue volumes. Based on Atropos results, GM volume decreased significantly (0.05% per year), while WM (0.03% per year) and CSF (0.02% per year) volumes significantly increase. This increase in WM volume is unlikely to be caused by any physical changes and might be due to the fact that Atropos tended to under-segment CSF. BISON estimates of the tissue volumes showed more modest changes in the volumes, with a slight decrease for GM (0.01% per year) and WM (0.01% per year), and an increase in the CSF (0.02% per year) volume.

An important concern when using a segmentation tool in aging and diseased populations is the presence of vascular pathology. WMHs of presumed vascular origins are very common findings in such cohorts (Wardlaw et al., 2015), and can be present in as high as 75% of the cases with dementia (van der Flier et al., 2018). Since they contribute to cognitive deficits and neurodegeneration, incorrectly segmenting them as GM can introduce a systematic error into studies in such populations. Our assessment of the tissue segmentations in 16 subjects with moderate to severe levels of WMHs showed that unlike Atropos, FAST from FSL, and SPM12, BISON provides accurate segmentations in presence of WMHs, making it a more reliable choice in populations that are likely to present with high loads of WMHs.

While generally validated on smaller datasets (N<20) of healthy young individuals, other automated tissue classification methods in the literature report similar or lower Dice Kappa values when compared with manual labels from real scans, e.g. 0.77-0.78 by Cocosco et al. (Cocosco et al., 2003), 0.83-0.84 by Van Leemput et al. (Van Leemput et al., 1999), 0.85 for GM and 0.86 for WM by Bazin et al. (Bazin and Pham, 2007), 0.80 for GM and 0.88 by Awate et al. (Awate et al., 2006), 0.4-0.8 by Ferreira da Silva (Ferreira da Silva, 2007), 0.78-0.84 for GM and 0.84-0.89 for WM by Tohka et al. (Tohka et al., 2010), 0.78 for GM, 0.85 for WM, and 0.22 for CSF by Greenspan et al. (Greenspan et al., 2006). This lower performance on real scans (compared to the synthetic data on which most methods are generally trained) is likely due to the added complexities caused by the heterogeneity in the real data; such as scanner or acquisition protocol differences, population differences, presence of pathology, and the inter-rater and intra-rater variabilities in the manually generated labels. In our experiments with the 20-Repeats dataset, the average test-retest Dice Kappa values were κ_GM =_ 0.92 ± 0.01, κ_WM =_ 0.91 ± 0.01, κ_CSF =_ 0.74 ± 0.03 for the manual segmentations, representing the level of variability in the manual segmentations that will inevitably lead to lower reported performances when automatic methods are compared against manual labels.

Since Neuromorphometrics dataset only includes T1-weighted images, the proposed method was trained and validated to segment tissue classes using only T1-weighted images. However, the pipeline has been developed to be able to handle (and the open-source software handles) any combination of other input sequences such as T2-weighted, PD-weighted and FLAIR images in addition to the T1-weighted images and can be retrained to segment other structures as well if a library manual labels is provided. The option of using additional FLAIR or T2-weighted images can be particularly useful in improving accuracies in presence of pathologies such as WMHs, tumors, stroke, etc.

Accurate and robust tissue classification from T1-weighted MR images is critical to many image processing and clinical applications. Due to the high variability of the tissue intensity profiles across different populations, image acquisition parameters, scanner models, manual segmentation protocols as well as rater variability, a tissue segmentation method that can provide robust classifications in different datasets is highly advantageous. Our results suggest that the proposed pipeline can provide accurate and robust tissue segmentations in multi-center and multi-scanner data, making it particularly useful for in analysing large multi-center MRI databases.

## Acknowledgement

We would like to acknowledge funding from the Famille Louise & André Charron. This work was also supported by grants from the Canadian Institutes of Health Research (MOP-111169), the Natural Sciences and Engineering Research Council of Canada (396395), les Fonds de Research Santé Québec Pfizer Innovation fund, the Levesque Foundation, the Douglas Hospital Research Centre and Foundation, the Government of Canada, and the Canada Fund for Innovation.

## References

Ad-Dab’bagh, Y., Lyttelton, O., Muehlboeck, J.S., Lepage, C., Einarson, D., Mok, K., Ivanov, O., Vincent, R.D., Lerch, J., Fombonne, E., others, 2006. The CIVET image-processing environment: a fully automated comprehensive pipeline for anatomical neuroimaging research, in: Proceedings of the 12th Annual Meeting of the Organization for Human Brain Mapping. Florence, Italy, p. 2266.

Aljabar, P., Heckemann, R.A., Hammers, A., Hajnal, J.V., Rueckert, D., 2009. Multi-atlas based segmentation of brain images: atlas selection and its effect on accuracy. Neuroimage 46, 726–738.

Aubert-Broche, B., Evans, A.C., Collins, L., 2006. A new improved version of the realistic digital brain phantom. NeuroImage 32, 138–145.

Avants, B.B., Tustison, N.J., Wu, J., Cook, P.A., Gee, J.C., 2011. An open source multivariate framework for n-tissue segmentation with evaluation on public data. Neuroinformatics 9, 381–400.

Awate, S.P., Tasdizen, T., Foster, N., Whitaker, R.T., 2006. Adaptive Markov modeling for mutual-information-based, unsupervised MRI brain-tissue classification. Med. Image Anal., The Eighth International Conference on Medical Imaging and Computer Assisted Intervention – MICCAI 2005 10, 726–739. https://doi.org/10.1016/j.media.2006.07.002

Bazin, P.-L., Pham, D.L., 2007. Topology-preserving tissue classification of magnetic resonance brain images. IEEE Trans. Med. Imaging 26, 487–496.

Beekly, D.L., Ramos, E.M., Van Belle, G., Deitrich, W., Clark, A.D., Jacka, M.E., Kukull, W.A., others, 2004. The National Alzheimer’s Coordinating Center (NACC) Database: an Alzheimer disease database. Alzheimer Dis. Assoc. Disord. 18, 270–277.

Breiman, L., 2001. Random Forests. Mach. Learn. 45, 5–32. https://doi.org/10.1023/A:1010933404324

Cabezas, M., Oliver, A., Lladó, X., Freixenet, J., Cuadra, M.B., 2011. A review of atlas-based segmentation for magnetic resonance brain images. Comput. Methods Programs Biomed. 104, e158–e177.

Caviness Jr, V.S., Lange, N.T., Makris, N., Herbert, M.R., Kennedy, D.N., 1999. MRI-based brain volumetrics: emergence of a developmental brain science. Brain Dev. 21, 289–295.

Cocosco, C.A., Zijdenbos, A.P., Evans, A.C., 2003. A fully automatic and robust brain MRI tissue classification method. Med. Image Anal., Medical Image Computing and Computer Assisted Intervention 7, 513–527. https://doi.org/10.1016/S1361-8415(03)00037-9

Collins, D.L., Evans, A.C., 1997. Animal: validation and applications of nonlinear registration-based segmentation. Int. J. Pattern Recognit. Artif. Intell. 11, 1271–1294.

Collins, D.L., Pruessner, J.C., 2010. Towards accurate, automatic segmentation of the hippocampus and amygdala from MRI by augmenting ANIMAL with a template library and label fusion. Neuroimage 52, 1355–1366.

Coupe, P., Yger, P., Prima, S., Hellier, P., Kervrann, C., Barillot, C., 2008. An Optimized Blockwise Nonlocal Means Denoising Filter for 3-D Magnetic Resonance Images. IEEE Trans. Med. Imaging 27, 425–441. https://doi.org/10.1109/TMI.2007.906087

Dadar, M., Fonov, V.S., Collins, D.L., Initiative, A.D.N., 2018a. A comparison of publicly available linear MRI stereotaxic registration techniques. NeuroImage 174, 191–200.

Dadar, M., Maranzano, J., Ducharme, S., Carmichael, O.T., Decarli, C., Collins, D.L., Initiative, A.D.N., 2018b. Validation of T 1w-based segmentations of white matter hyperintensity volumes in large-scale datasets of aging. Hum. Brain Mapp. 39, 1093–1107.

Dadar, M., Maranzano, J., Ducharme, S., Collins, D.L., 2019. White matter in different regions evolves differently during progression to dementia. Neurobiol. Aging 76, 71–79. https://doi.org/10.1016/j.neurobiolaging.2018.12.004

Dadar, M., Maranzano, J., Misquitta, K., Anor, C.J., Fonov, V.S., Tartaglia, M.C., Carmichael, O.T., Decarli, C., Collins, D.L., Alzheimer’s Disease Neuroimaging Initiative, 2017a. Performance comparison of 10 different classification techniques in segmenting white matter hyperintensities in aging. NeuroImage 157, 233–249. https://doi.org/10.1016/j.neuroimage.2017.06.009

Dadar, M., Pascoal, T., Manitsirikul, S., Misquitta, K., Tartaglia, C., Brietner, J., Rosa-Neto, P., Carmichael, O., DeCarli, C., Collins, D.L., 2017b. Validation of a Regression Technique for Segmentation of White Matter Hyperintensities in Alzheimer’s Disease. IEEE Trans. Med. Imaging.

Dadar, M., Zeighami, Y., Yau, Y., Fereshtehnejad, S.-M., Maranzano, J., Postuma, R.B., Dagher, A., Collins, D.L., 2018c. White matter hyperintensities are linked to future cognitive decline in de novo Parkinson’s disease patients. NeuroImage Clin. 20, 892–900.

De Boer, R., Vrooman, H.A., Van Der Lijn, F., Vernooij, M.W., Ikram, M.A., Van Der Lugt, A., Breteler, M.M., Niessen, W.J., 2009. White matter lesion extension to automatic brain tissue segmentation on MRI. Neuroimage 45, 1151–1161.

Dice, L.R., 1945. Measures of the amount of ecologic association between species. Ecology 26, 297–302.

Duda, R.O., Hart, P.E., Stork, D.G., 2001. Pattern Classification, A Wiley-Interscience Publication John Wiley & Sons. Inc.

Ferreira da Silva, A.R., 2007. A Dirichlet process mixture model for brain MRI tissue classification. Med. Image Anal. 11, 169–182. https://doi.org/10.1016/j.media.2006.12.002

Fischl, B., Salat, D.H., Busa, E., Albert, M., Dieterich, M., Haselgrove, C., Van Der Kouwe, A., Killiany, R., Kennedy, D., Klaveness, S., others, 2002. Whole brain segmentation: automated labeling of neuroanatomical structures in the human brain. Neuron 33, 341–355.

González-Villà, S., Oliver, A., Valverde, S., Wang, L., Zwiggelaar, R., Lladó, X., 2016. A review on brain structures segmentation in magnetic resonance imaging. Artif. Intell. Med. 73, 45–69. https://doi.org/10.1016/j.artmed.2016.09.001

Greenspan, H., Ruf, A., Goldberger, J., 2006. Constrained Gaussian mixture model framework for automatic segmentation of MR brain images. IEEE Trans. Med. Imaging 25, 1233–1245. https://doi.org/10.1109/TMI.2006.880668

Hinton, L., Carter, K., Reed, B.R., Beckett, L., Lara, E., DeCarli, C., Mungas, D., 2010. Recruitment of a community-based cohort for research on diversity and risk of dementia. Alzheimer Dis. Assoc. Disord. 24, 234.

Jo, H.J., Saad, Z.S., Simmons, W.K., Milbury, L.A., Cox, R.W., 2010. Mapping sources of correlation in resting state FMRI, with artifact detection and removal. Neuroimage 52, 571–582.

Klein, A., Tourville, J., 2012. 101 labeled brain images and a consistent human cortical labeling protocol. Front. Neurosci. 6, 171.

Lötjönen, J.M., Wolz, R., Koikkalainen, J.R., Thurfjell, L., Waldemar, G., Soininen, H., Rueckert, D., Initiative, A.D.N., 2010. Fast and robust multi-atlas segmentation of brain magnetic resonance images. Neuroimage 49, 2352–2365.

Makropoulos, A., Gousias, I.S., Ledig, C., Aljabar, P., Serag, A., Hajnal, J.V., Edwards, A.D., Counsell, S.J., Rueckert, D., 2014. Automatic whole brain MRI segmentation of the developing neonatal brain. IEEE Trans. Med. Imaging 33, 1818–1831.

Marek, K., Jennings, D., Lasch, S., Siderowf, A., Tanner, C., Simuni, T., Coffey, C., Kieburtz, K., Flagg, E., Chowdhury, S., Poewe, W., Mollenhauer, B., Klinik, P.-E., Sherer, T., Frasier, M., Meunier, C., Rudolph, A., Casaceli, C., Seibyl, J., Mendick, S., Schuff, N., Zhang, Y., Toga, A., Crawford, K., Ansbach, A., De Blasio, P., Piovella, M., Trojanowski, J., Shaw, L., Singleton, A., Hawkins, K., Eberling, J., Brooks, Deborah, Russell, D., Leary, L., Factor, S., Sommerfeld, B., Hogarth, P., Pighetti, E., Williams, K., Standaert, D., Guthrie, S., Hauser, R., Delgado, H., Jankovic, J., Hunter, C., Stern, M., Tran, B., Leverenz, J., Baca, M., Frank, S., Thomas, C.-A., Richard, I., Deeley, C., Rees, L., Sprenger, F., Lang, E., Shill, H., Obradov, S., Fernandez, H., Winters, A., Berg, D., Gauss, K., Galasko, D., Fontaine, D., Mari, Z., Gerstenhaber, M., Brooks, David, Malloy, S., Barone, P., Longo, K., Comery, T., Ravina, B., Grachev, I., Gallagher, K., Collins, M., Widnell, K.L., Ostrowizki, S., Fontoura, P., Ho, T., Luthman, J., Brug, M. van der, Reith, A.D., Taylor, P., 2011. The Parkinson Progression Marker Initiative (PPMI). Prog. Neurobiol., Biological Markers for Neurodegenerative Diseases 95, 629–635. https://doi.org/10.1016/j.pneurobio.2011.09.005

Mateos-Pérez, J.M., Dadar, M., Lacalle-Aurioles, M., Iturria-Medina, Y., Zeighami, Y., Evans, A.C., 2018. Structural neuroimaging as clinical predictor: A review of machine learning applications. NeuroImage Clin. https://doi.org/10.1016/j.nicl.2018.08.019

Pedregosa, F., Varoquaux, G., Gramfort, A., Michel, V., Thirion, B., Grisel, O., Blondel, M., Prettenhofer, P., Weiss, R., Dubourg, V., others, 2011. Scikit-learn: Machine learning in Python. J. Mach. Learn. Res. 12, 2825–2830.

Penny, W.D., Friston, K.J., Ashburner, J.T., Kiebel, S.J., Nichols, T.E., 2011. Statistical parametric mapping: the analysis of functional brain images. Academic press.

Sajja, B.R., Datta, S., He, R., Mehta, M., Gupta, R.K., Wolinsky, J.S., Narayana, P.A., 2006. Unified approach for multiple sclerosis lesion segmentation on brain MRI. Ann. Biomed. Eng. 34, 142–151.

Scherrer, B., Forbes, F., Garbay, C., Dojat, M., 2009. Distributed local MRF models for tissue and structure brain segmentation. IEEE Trans. Med. Imaging 28, 1278–1295.

Schmidt, P., Gaser, C., Arsic, M., Buck, D., Förschler, A., Berthele, A., Hoshi, M., Ilg, R., Schmid, V.J., Zimmer, C., others, 2012. An automated tool for detection of FLAIR-hyperintense white-matter lesions in multiple sclerosis. Neuroimage 59, 3774–3783.

Shen, D., Davatzikos, C., 2001. HAMMER: hierarchical attribute matching mechanism for elastic registration, in: Proceedings IEEE Workshop on Mathematical Methods in Biomedical Image Analysis (MMBIA 2001). IEEE, pp. 29–36.

Simões, R., Mönninghoff, C., Dlugaj, M., Weimar, C., Wanke, I., van Walsum, A.-M. van C., Slump, C., 2013. Automatic segmentation of cerebral white matter hyperintensities using only 3D FLAIR images. Magn. Reson. Imaging 31, 1182–1189.

Sled, J.G., Zijdenbos, A.P., Evans, A.C., 1998. A nonparametric method for automatic correction of intensity nonuniformity in MRI data. Med. Imaging IEEE Trans. On 17, 87–97.

Steenwijk, M.D., Pouwels, P.J., Daams, M., van Dalen, J.W., Caan, M.W., Richard, E., Barkhof, F., Vrenken, H., 2013. Accurate white matter lesion segmentation by k nearest neighbor classification with tissue type priors (kNN-TTPs). NeuroImage Clin. 3, 462–469.

Tohka, J., Dinov, I.D., Shattuck, D.W., Toga, A.W., 2010. Brain MRI tissue classification based on local Markov random fields. Magn. Reson. Imaging 28, 557–573. https://doi.org/10.1016/j.mri.2009.12.012

Tremblay-Mercier, J., Madjar, C., Etienne, P., Poirier, J., Breitner, J., 2014. A PROGRAM OF PRE-SYMPTOMATIC EVALUATION OF EXPERIMENTAL OR NOVEL TREATMENTS FOR ALZHEIMER’S DISEASE (PREVENT-AD): DESIGN, METHODS, AND PERSPECTIVES. Alzheimers Dement. J. Alzheimers Assoc. 10, P808.

van der Flier, W.M., Skoog, I., Schneider, J.A., Pantoni, L., Mok, V., Chen, C.L.H., Scheltens, P., 2018. Vascular cognitive impairment. Nat. Rev. Dis. Primer 4, 18003. https://doi.org/10.1038/nrdp.2018.3

Van Leemput, K., Maes, F., Vandermeulen, D., Suetens, P., 1999. Automated model-based tissue classification of MR images of the brain. IEEE Trans. Med. Imaging 18, 897–908.

Wardlaw, J.M., Hernández, M.C.V., Muñoz-Maniega, S., 2015. What are white matter hyperintensities made of? Relevance to vascular cognitive impairment. J. Am. Heart Assoc. 4, e001140.

Worth, A.J., Makris, N., Kennedy, D.N., Caviness Jr, V.S., 2001. Accountability in methodology and analysis for clinical trials involving quantitative measurements of MR brain images. Technical Report TR20011117, Neuromorphometrics, Inc.

Wu, M., Rosano, C., Lopez-Garcia, P., Carter, C.S., Aizenstein, H.J., 2007. Optimum template selection for atlas-based segmentation. NeuroImage 34, 1612–1618.

Zhang, Y., Brady, M., Smith, S., 2001. Segmentation of brain MR images through a hidden Markov random field model and the expectation-maximization algorithm. IEEE Trans. Med. Imaging 20, 45–57.

